# Patterns of grass (Poaceae) species distribution and richness across India

**DOI:** 10.1101/2025.05.06.651586

**Authors:** Mukta Mande, Atul A. Joshi, Harinandanan Paramjyothi, Jayashree Ratnam, Mahesh Sankaran

**Author notes:** A.A.J.: Department of Physical and Natural Sciences, FLAME University, Pune 412115, Maharashtra, India. H.P.: Research Institute for the Environment and Livelihoods, Charles Darwin University Darwin NT 0909, Australia.

## Abstract

Tropical grassy biomes (TGBs) cover a substantial portion of Earth’s land surface area and support a rich diversity of fauna and flora. Yet, despite their importance for biodiversity and human livelihoods, the TGBs of South Asia, including India, have historically been undervalued, and as result, understudied. Here, we address this gap with the first subcontinental-scale analyses of grass species richness and distribution across the Indian region. Using data on species occurrences collated from multiple regional floras, we describe the distribution of families, functional groups and species of Poaecea across this region, and their variation as a function of climate and geomorphology. India hosts >1100 grass species, comprising ∼10% of global grass richness. Over half of these were perennial C4 grasses, with Panicoideae being the most speciose subfamily in the country. Consistent with patterns observed globally, the PACMAD clade consisting of both C3 and C4 species was more speciose in warmer regions, while the C3 subfamilies of the BEP clade, Pooideae and Bambusoideae, were more abundant in cooler regions of the country. Both total and C4 species richness were greatest in warmer districts with high moisture availability, while cooler and wetter districts with low precipitation seasonality supported greater C3 species richness. Our results spotlight previously unrecognized areas of high grass species richness in the arid and semi-arid areas of North-West India and the Deccan Peninsula in India, and call for conservation attention to neglected regions that are outside known biodiversity hotspots and protected areas.

## 1. Introduction

Tropical grassy biomes (TGBs), ranging from open grasslands to savannas, dominate terrestrial regions of the tropics (Parr et al., 2014). They are characterised by a continuous C_4_ grassy understory with discontinuous woody strata (Scholes & Archer, 1997). TGBs cover ∼40% of Earth’s land surface area and provide critical services in the form of grazing lands, fuel and food for nearly a fifth of the world’s human population. Globally, there are more than 11,000 species of grasses across all biomes, but while these constitute less than 5% of global plant diversity, they account for nearly 25% of global net primary production (Beer et al., 2010; Linder et al., 2018; Soreng et al., 2022). Grasslands also store an estimated 10-30% of global soil carbon (Gibson, 2009; Grace et al., 2006; Lehmann & Parr, 2016; Parr et al., 2014). However, despite supporting diverse and abundant assemblages of extant mammalian megafauna, and a rich diversity of other mammals, birds, amphibians and vascular plants the factors that structure the diversity of TGBs remains poorly documented as compared to tropical forests (Murphy et al., 2016; Sankaran & Ratnam, 2013). Given the high levels of threats these biomes face globally, particularly in terms of land conversion to other uses, understanding the factors that regulate patterns of species diversity and distribution in these biomes is critical for their effective conservation (Murphy et al., 2016).

Although several earlier studies have investigated patterns of faunal and floral diversity of grassy biomes across the Americas, Neotropics, Afro-tropics, Australia and China (Bocksberger et al., 2016; Bremond et al., 2012; Griffith et al., 2015; Hattersley, 1983; Liu et al., 2009; Teeri & Stowe, 1976), the TGBs of South Asia, including India, have remained largely unrecognized (Murphy et al., 2016; Ratnam et al., 2016). Colonial-era policies that gave primacy to timber-bearing forests and classified sparsely-treed, open grassy biomes as “wastelands” have left a legacy of TGB’s being undervalued, converted to other land uses and largely missed from the scientific discourse of the region, until recently (Lahiri & Reddy, 2025; Ratnam et al., 2016; Tölgyesi et al., 2024; Vanak et al., 2014). These recent studies show that TGBs currently cover around 10% of the land surface area of India (Madhusudan & Vanak, 2022), host a substantial diversity of endemic plants, including grasses (Nerlekar et al., 2022; Sankaran, 2009), and support the livelihoods of millions of marginalized pastoral people (Sheth et al., in press). However, a regional-scale understanding of the vegetation communities in these biomes, which are shaped primarily by the grass species that dominate in these ecosystems and define their productivity, diversity and functional ecology is still missing. Given their defining role in the ecology of TGBs, understanding the factors regulating the distribution of grasses is key to predicting their future dynamics under scenarios of global change (Bocksberger et al., 2016).

In this study, we use species occurrence records, collated at the district level from published floras, to examine patterns of grass species richness and distribution across broad-scale environmental gradients in the Indian subcontinent. Due to its unique biogeography, the Indian subcontinent hosts a diverse array of biomes spanning a range of temperature, seasonality, altitude and precipitation regimes. Poaceae (grasses) dominate the understory of many of these biomes. Here, we examine how patterns of total species richness, rarity, and richness of different grass functional groups (C_3_ and C_4_) and major lineages of the Poaceae change across the Indian subcontinent, and across different biogeographic zones in the country, and evaluate the role of climatic factors in driving these patterns.

Overall, we expected grass species richness to be highest in TGBs (open ecosystems such as savannas and grasslands where grasses dominate the understory) and lowest in forested ecosystems with closed canopies, and grass species richness to generally increase with increasing topographic complexity of the district. Further, we expected C_4_ grass species richness to be greatest in warm and arid regions i.e., areas with high mean annual temperatures, high growing season temperatures and low to intermediate rainfall with high seasonality, and C_3_ richness to be highest in cooler and wetter regions with low seasonality in rainfall (Edwards & Still, 2008; Griffith et al., 2015; Murphy & Bowman, 2007; Teeri & Stowe, 1976; Tieszen et al., 1979). C_4_ plants have a competitive advantage over C_3_ plants at high temperatures because they have evolved mechanisms to reduce the negative effects of photorespiration - a major source of carbon loss during photosynthesis, which imposes major limitations to photosynthesis and growth of C_3_ plants at high temperatures (Christin & Osborne, 2014; Edwards & Still, 2008; Ehleringer et al., 1997; Griffith et al., 2015; Sage, 2004; Still et al., 2003; Taylor et al., 2010, 2014). C_4_ plants also have a competitive advantage over C_3_ plants under more arid conditions as their CO_2_ concentrating mechanism allows them to have lower stomatal conductance and reduce water loss (Edwards & Still, 2008; Sage, 2004). Lastly, C4 abundance has been linked to rainfall seasonality, specifically warm season rainfall in Australia and Western United States (Havrilla et al., 2023; Murphy & Bowman, 2007).

We also assessed how the richness and distribution of different grass lineages - the ‘BEP’ and ‘PACMAD’ clades - differed between different biomes and climate regimes. We expected the BEP clade (or BOP clade; Bambusoideae, Oryzoideae (syn. Ehrhartoideae) and Pooideae), which comprises ∼3300 species globally including the bamboos, relatives of rice and Pooideae that radiated extensively in cold and temperate regions (Edwards & Still, 2008) to dominate in cooler areas at higher elevations and latitudes, and in wetter areas (as in the case of bamboos) of the subcontinent. On the other hand, we expected the ‘PACMAD’ clade (Panicoideae, Arundinoideae, Chloridoideae, Micrairoideae, Aristidoideae, and Danthonioideae) which contains all C_4_ grasses to dominate in open habitats in warmer regions of the subcontinent (Edwards & Still, 2008).

## 2. Methods

### 2.1 Dataset

The dataset used for this study was compiled from various regional floras published from 1966 to 2011 (Table S1). The dataset includes the list of grass species present in each district of India. Additionally, we also recorded taxonomic information (subfamily, super-tribe, tribe, sub-tribe, subspecies and variety), life history, growth form, photosynthetic pathway (C_3_ vs C_4_), origin, habitat and phenology where reported in the floras. For species where information on photosynthetic pathway was not reported in the flora, data were taken from Osborne et al., (2014) (Osborne et al., 2014). All species belonging to the BEP clade were assigned the C_3_ pathway. Species names were updated using the ‘*WorldFlora*’ package in R as per the taxonomic backbone static version dated March 2023. Instances with multiple matches were manually corrected case by case.

The district names in the collated dataset from the floras were corrected as per the 2011 census list from the online maps portal of Survey of India. The shape file used for analysis was updated as per this list of corrected district names.

### 2.2 Analysis

We evaluated the role of multiple climatic factors in determining overall grass species richness, C_3_ vs C_4_ richness, and the richness of the dominant clades (Fig. S1-4) at the district level across India. Our predictor variables included measures of moisture availability and temperature, both of which have been shown to be important drivers of grass functional group richness (see introduction). In addition, we also included district area and measures of topographic and climatic heterogeneity at the district level as predictors, given their importance in regulating taxonomic and functional richness of grasses (Liu et al., 2009). Our final list of predictors included mean annual precipitation (MAP, mm), moisture index (MI or aridity index, calculated as MAP/ Potential Evapotranspiration), precipitation seasonality (PS, coefficient of variation in monthly rainfall), mean annual temperature (MAT, °C), maximum temperature of wettest quarter (TW), mean annual precipitation range of the district (MAP range), moisture index range of the district (MI range), mean annual temperature range of the district (MAT range), a topographic factor - elevational range of each district, and district area.

Climatic data, MAP, MAT, PS, MAP range and MAT range were extracted from WorldClim 2.1 at 30-second resolution (∼1 km^2^ at equator) (Fick & Hijmans, 2017) and TW was extracted from CHIRTSmonthly at 0.05° resolution (∼5.5 km^2^ at equator) (Funk et al., 2015) and averaged to get a mean value for each district. We chose to use maximum, rather than mean, temperature of the wettest month since the former better represents day-time growing temperatures as compared to mean temperatures (Griffith et al., 2015). Data on the moisture index (MI and MI range) were extracted from the CGIAR Global Aridity Index and Potential Evapotranspiration database ver. 2.0 (Trabucco & Zomer, 2019), and similarly averaged across all pixels in each district. We used MI range and MAT range of a district as a proxy for the variation in climatic factors in the district. Elevation data were extracted from WorldClim 2.1 at 2.5 minute resolution (Fick & Hijmans, 2017). District-level means and ranges for all variables were calculated using the zonal statistics plugin in QGIS 3.22.6 (QGIS Development Team, n.d.).

Each district in our dataset was assigned a biogeographic zone based on the biogeographic classification of the Indian subcontinent by Rodgers et al. (2002) (Rodgers et al., 2002) (Fig. 1). In cases where the district spanned multiple zones, the district was assigned the biogeographic zone that covered the maximum area of the district. The number of biogeographic zones in each district was also noted. For each biogeographic zone, we calculated the total number of grass species, richness of different functional groups (C_3_ vs C_4_) and subfamilies of Poaceae, and life history strategies (annual vs perennial) by summing across all districts assigned to that biogeographic zone. We also examined patterns of grass species range size distributions (widespread versus geographically restricted) by calculating the number of different districts from which each species was reported.

**Figure 1:**
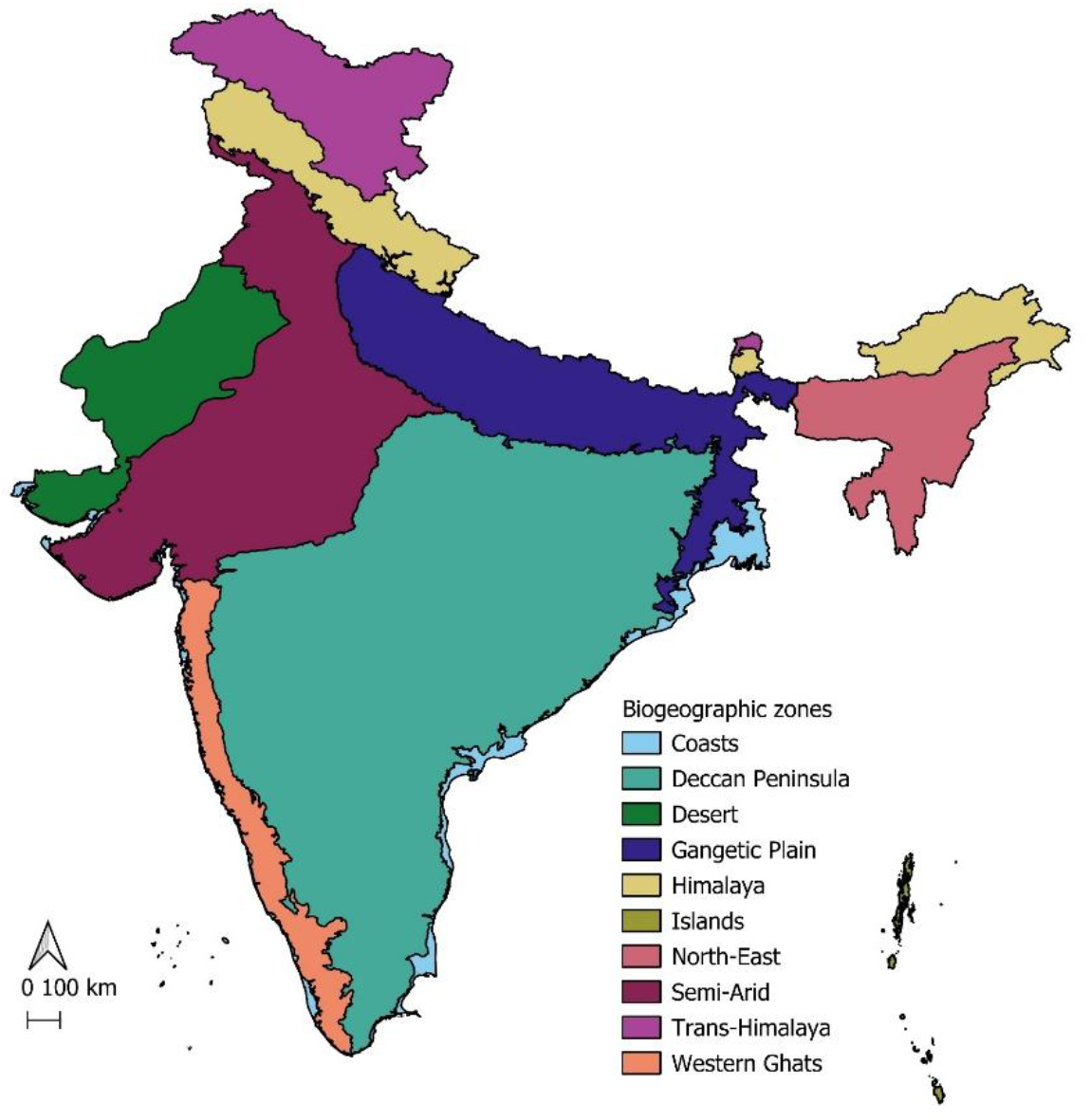
Map showing the biogeographic zones of India as per Rodgers et al. (2002)

Because our different indices of water availability and temperature were correlated, we ran multiple models with different combinations of predictor variables (e.g. MAP and MAP range vs MI and MI range, MAT vs TW; see Table S2). Further, because elevational range and MAT range were both highly correlated with mean annual temperature of the district (MAT; r = -0.83 in both cases), they were both dropped from all further analyses. We used generalized linear models with a negative binomial error distribution and log link function to evaluate the effects of climatic predictors on total richness, C_3_ and C_4_ species richness, and richness of different sub-families. We used generalized linear models with a binomial error distribution and logit link to check for the effect of different climatic predictors on the proportion of C_4_ species in districts. We included log (area of the district) as a covariate in all models to account for the differences in area between districts. Two biogeographic zones - the Trans-Himalayas and Islands - were excluded from the analysis due to insufficient data. We only report results from the best-performing models (lowest AIC) here.

All analyses were carried out using R 4.3.0 (R Core Team, 2023) and RStudio 2023.03.1 (R Core Team, 2023). We used the *tidyverse* package (Wickham et al., 2019) for all data processing and the *sjPlot* package for data visualization (Lüdecke, 2024).

## 3. Results

### 3.1 Species distribution patterns in the Indian subcontinent

Our final dataset describes the distribution patterns of 1121 grass species belonging to 249 genera (of the 768 described thus far globally; (Soreng et al., 2022)) and 10 of the 12 subfamilies of Poaceae. Of these, 652 were C_4_, 430 were C_3_, 12 were either C_3_ or C_4_, and 27 had unkown photosynthetic pathway.

Northeast India, the northern and southern Western Ghats, and the north-western arid zone were hotspots of total grass species richness in India (Fig. 2a). These same areas, with the addition of the Deccan plateau, were also the centers of C_4_ species richness in the country (Fig. 2b). On the other hand, C3 species richness was highest in the Northeast and the western Himalaya (Fig. 2c).

**Figure 2:**
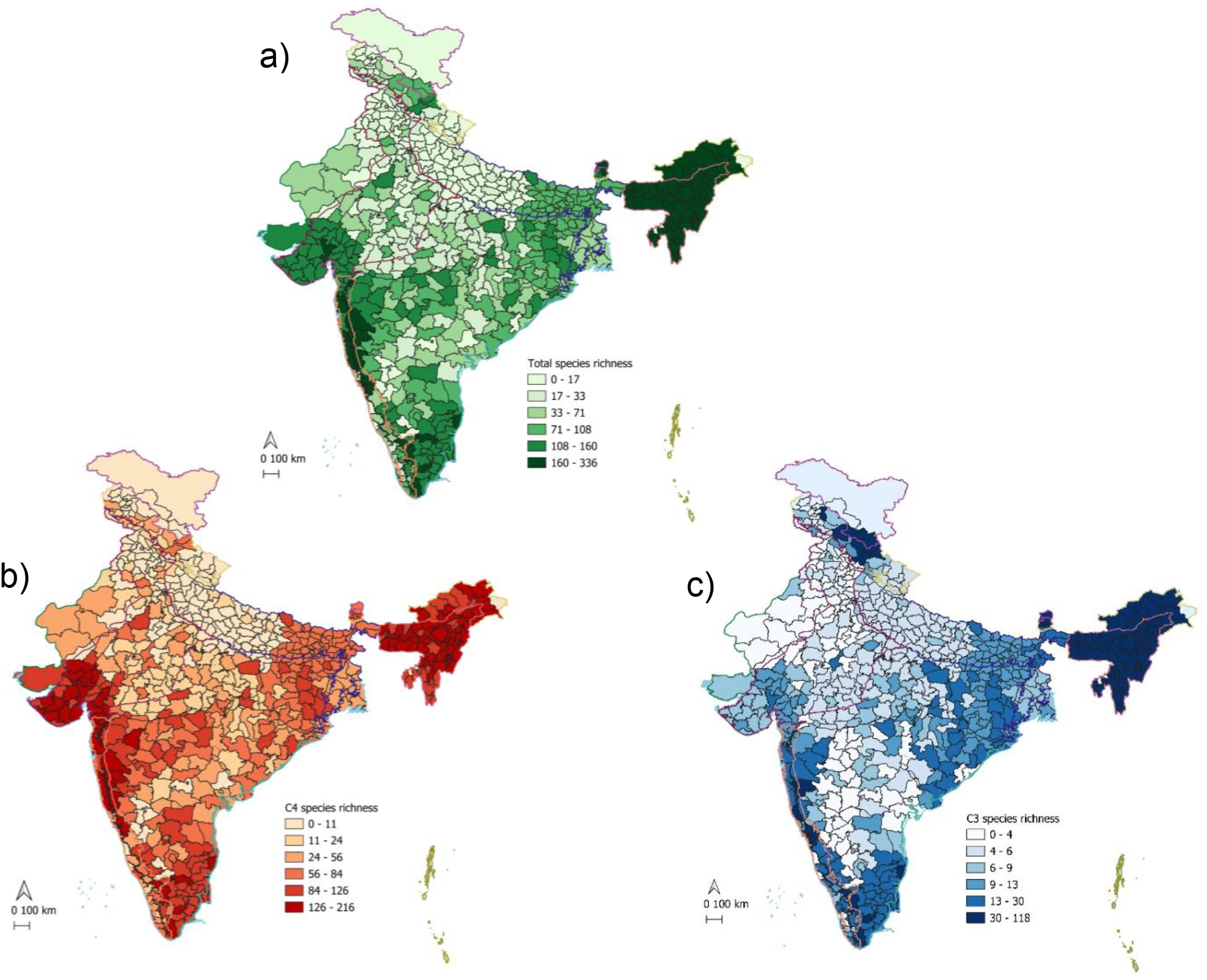
a) Total species richness of Poaceae at district level across India, b) C4 grass species richness at district level across India, c) C3 grass species richness at district level across India

Amongst the biogeographic zones, the Deccan peninsula had the highest species richness (622) followed by the Western Ghats (535) and Himalayas (524), while the Islands had the lowest species richness with 37 species (Fig. 3a). On the other hand, species richness per unit area was highest in the Islands (0.0051 species/km^2^) whereas the Deccan peninsula had the lowest number of species per unit area (0.0005 species/km^2^) (Fig. 3b).

**Figure 3:**
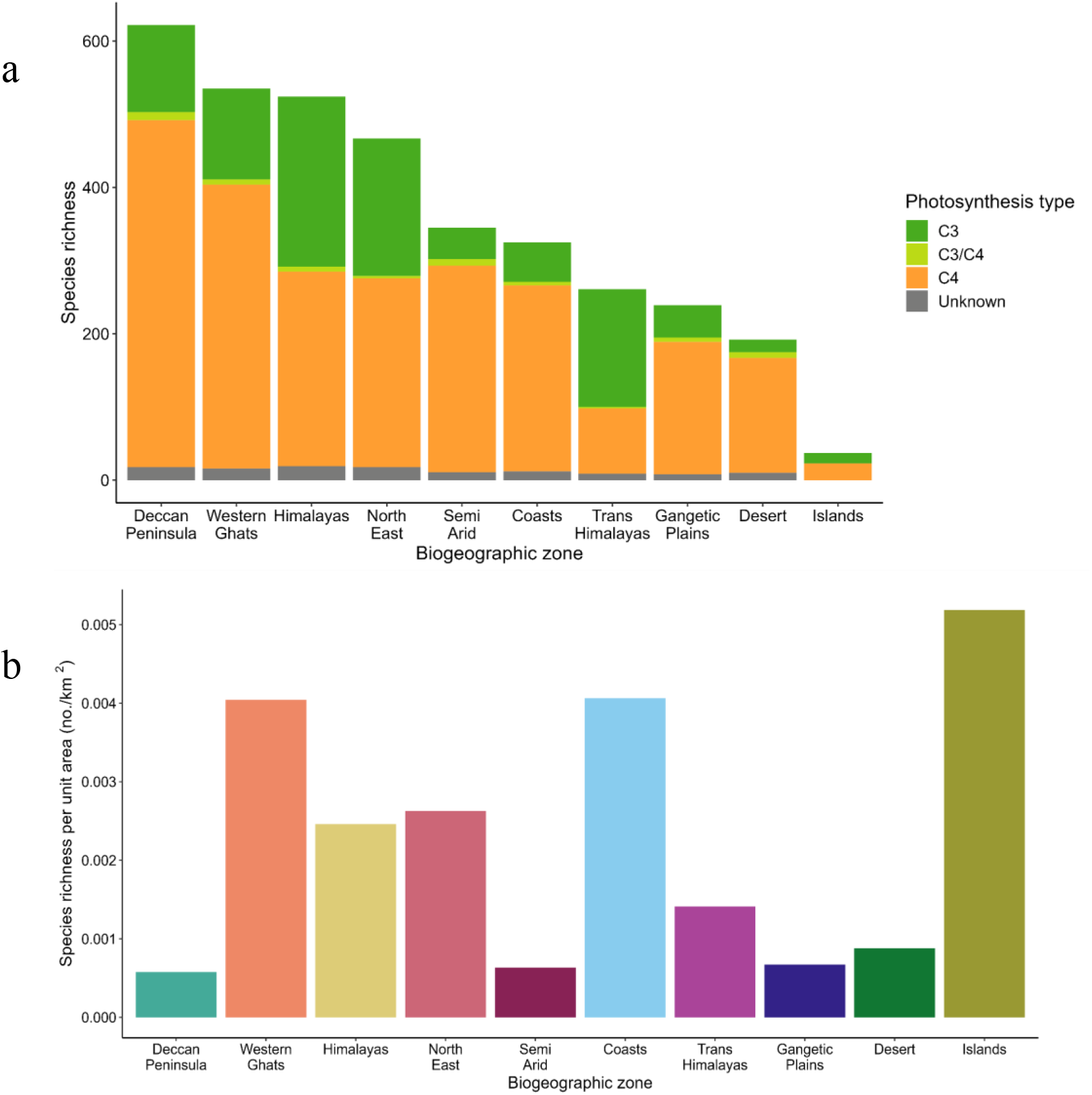
a) Total species richness and richness of different photosynthetic types in different biogeographic ones, and b) Species richness per unit area in different biogeographic zones.

All biogeographic zones, except the Trans-Himalaya, supported more C_4_ grasses than C_3_ grasses (Fig. 3a). C_4_ species richness was highest in the Deccan Peninsula with 474 species followed by the Western Ghats with 388 species. In contrast, the richness of C_3_ species was highest in the Himalayas (232) followed by the North-East (188) (Fig. 3a).

Most grass species were range-restricted, i.e. occurred in just one or a few districts (Fig 4a). A total of 206 species were recorded in just a single district each, while only 4 species occurred in more than 350 districts (Fig. 4a). The Western Ghats had the highest number - 48 - of range-restricted species (i.e. recorded from just 1 district), followed by the Himalayas with 42 species. The Gangetic Plains on the other hand had only one such species (Fig. 4b).

**Figure 4:**
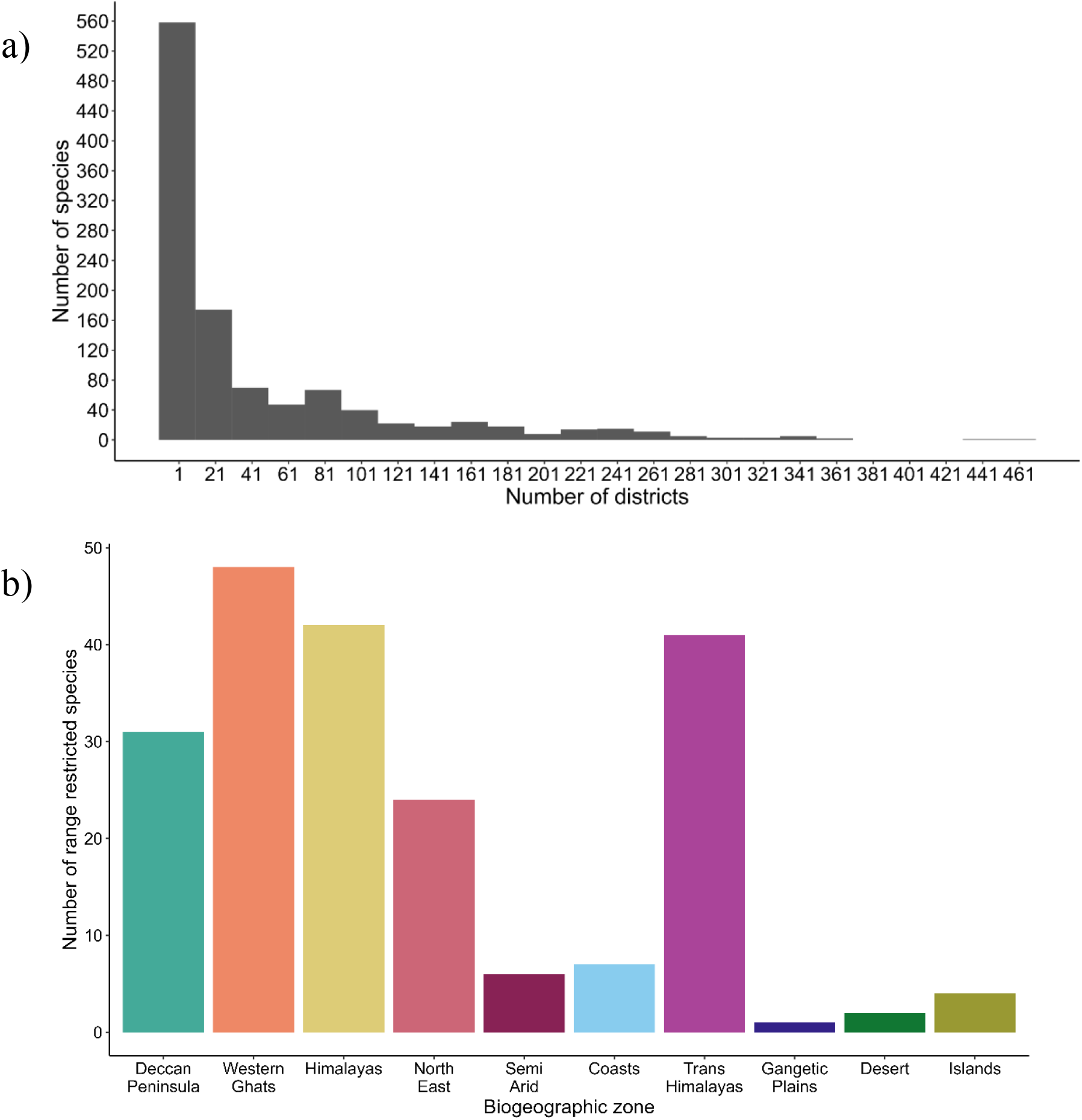
a) Number of species occurring in different number of districts and b) Number of species with distribution restricted to one district in different biogeographic zones

At the country scale, Panicoideae was the most speciose subfamily with 537 species, followed by Pooideae with 232 species (Fig. 5b). Panicoideae was also the most species-rich subfamily across all biogeographic zones (Fig. 5a) except the Himalayas and Trans-Himalayas where Pooideae dominated, and the Desert where Chloridoideae were most speciose (Fig. 5a).

**Figure 5:**
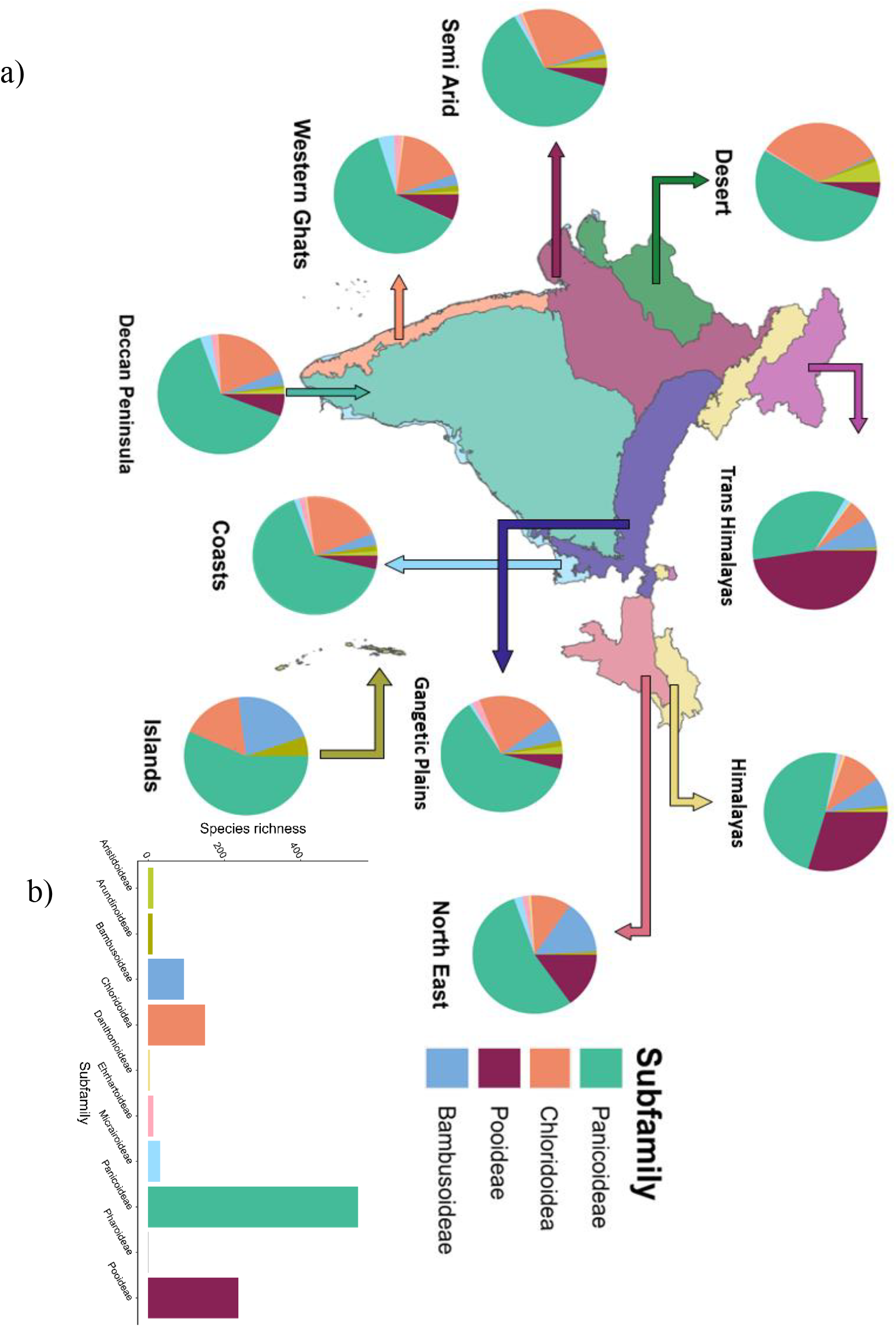
a) Species richness of different subfamilies of Poaceae across the Indian subcontinent, b) Species richness of all Poaceae subfamilies in different biogeographic zones

Most of the grass species documented (713) were perennials. Perennials accounted for half or more of the species in all biogeographic zones (Fig. 6). Although the number of annuals (255) was the highest in the Deccan peninsula, the proportion of annual species was highest in the Desert zone closely followed by the Semi-arid zone.

**Figure 6:**
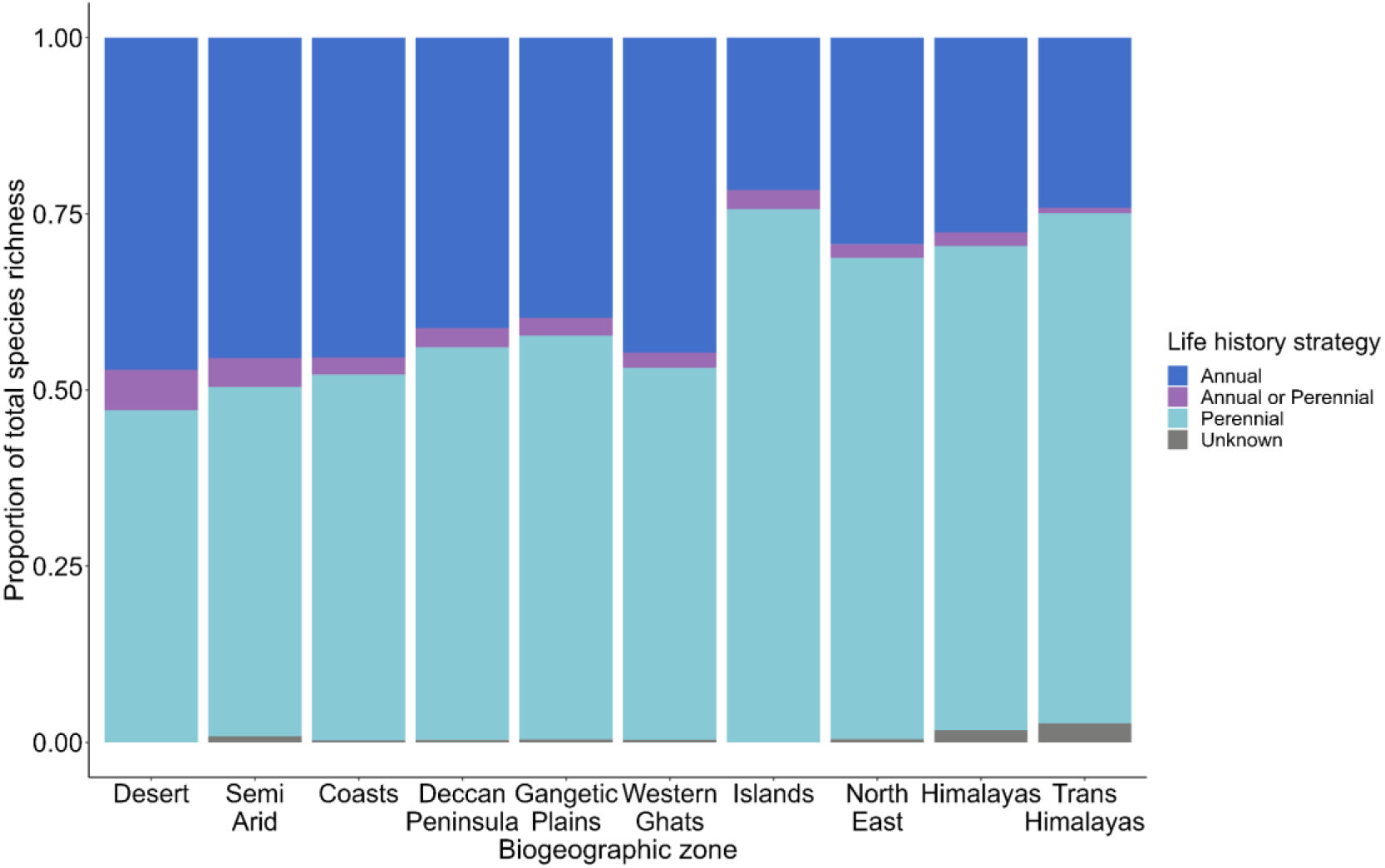
Proportion of annual and perennial grass species in different biogeographic zones of India.

### 3.2 Relationship of species richness with climate

Our final models explained 25%, 18%, and 64% of the variation in total, C_4_ and C_3_ species richness, respectively (Table S3, S4, S5). Not surprisingly, area was an important predictor of total, C4 and C3 grass species richness as well as richness of different subfamilies in a district (Fig. 7, Fig. S1, S2, S3, S4, Table S3, S4, S5). Both total and C4 species richness were greater in warmer districts with high moisture availability, wide MI ranges and low precipitation seasonality (Fig. 7, Table S3, S4). On the other hand, districts which were cooler during the wettest quarter with high moisture availability and lower seasonality of precipitation supported greater C3 species richness (Fig. 7). The highest proportion of C4 species was found in arid districts with high precipitation seasonality (Table S6, Fig. 8).

**Figure 7:**
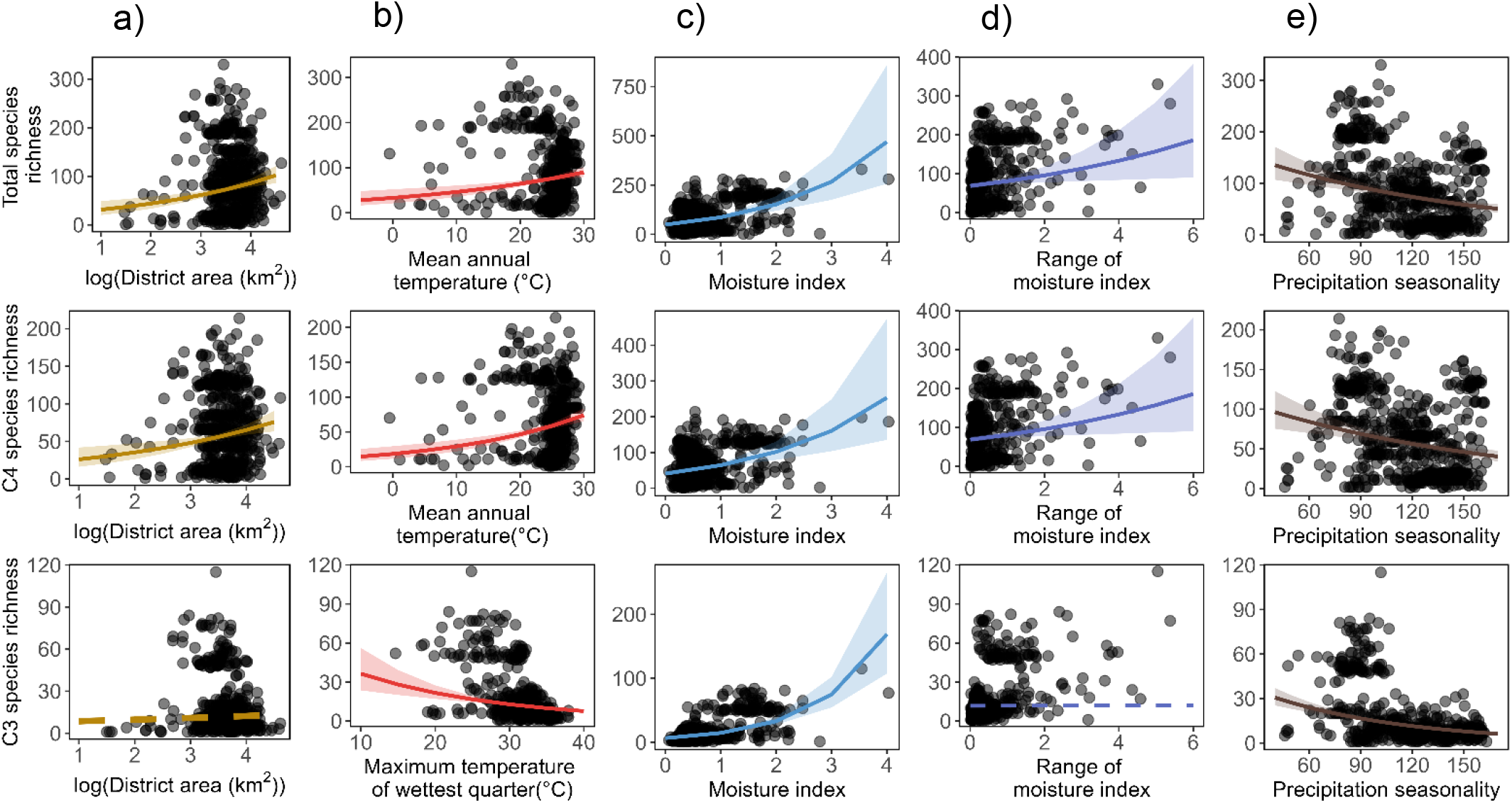
Relationship of corrected total, C4 and C3 species richness with: a) log(District area), b) Annual mean temperature (AMT), c) Moisture index, d) Moisture index range and e) Precipitation seasonality. The lines indicate model predictions and ribbons indicate 95% confidence intervals, dashed lines represent non-significant relationships.

**Figure 8:**
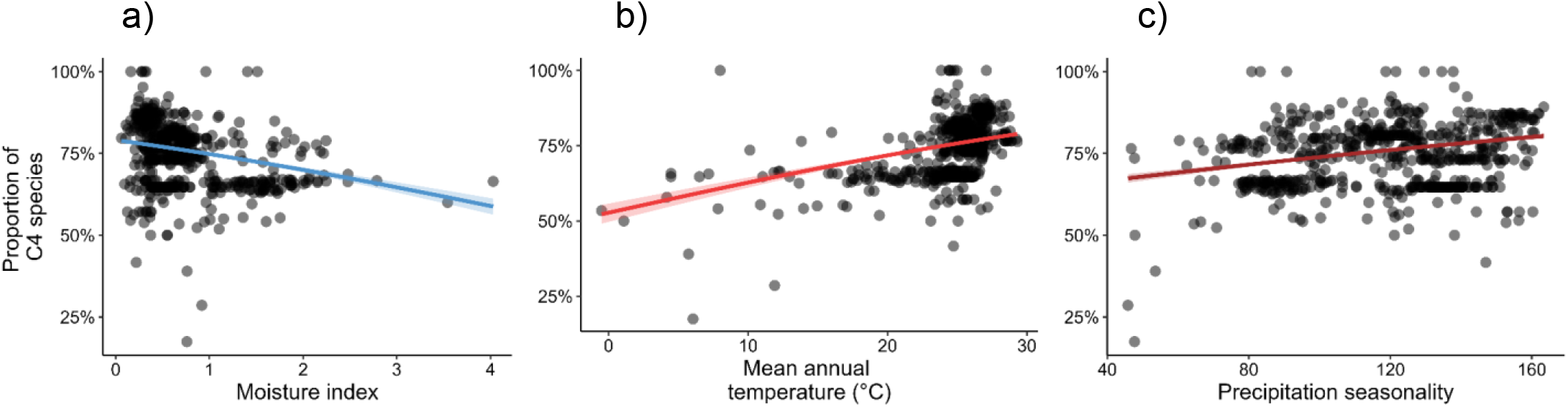
Relationship of the proportion of C4 species with: a) Moisture index, b) Annual mean temperature, c) Precipitation seasonality. The line indicates model predictions and ribbon indicate 95% confidence intervals; dashed line represents non-significant relationships.

In terms of the different subfamilies, the drivers of climate-richness patterns of Panicoideae were similar to those of total species richness (Fig.S1). Chloridoideae richness, on the other hand, was unrelated with MI. Chloridoideae richness was high in warmer districts with a wide range of MI values and low seasonality in precipitation (Fig.S2). The C3 subfamily of Pooideae was most species rich in cool districts with low precipitation seasonality and wide variation in MI values (Fig.S3). Similarly, richness of Bambusoideae was high in cooler and wetter districts with a wide range of MI values and low precipitation seasonality (Fig.S4).

## 4. Discussion

The tropical grassy biomes of the Indian subcontinent, cover around 10% of its land surface area (Madhusudan & Vanak, 2022), but have received scant scientific and conservation attention. Our collation of data from across this region shows that these TGBs host a substantial diversity of grasses (>1100 species) comprising nearly 10% of the world’s grass species and more than 30% of the world’s grass genera (Kellogg et al., 2020; Soreng et al., 2022). The majority of these (> 60%) are perennial warm-season (C4) grasses. Panicoideae is the most species-rich subfamily, constituting nearly half of all grass species in the subcontinent. Both total and C4 grass species richness appear to be concentrated in two hotspots - one in the east of the country, coinciding with the Indo-Burma biodiversity hotspot with high topographic heterogeneity, and another in the west, which includes the northern portion of the Western Ghats biodiversity hotspot, and notably, parts of the arid northwest (Fig 2a, b). Additionally, many districts of the Deccan Peninsula, which like the arid northwest, is generally not recognized for diversity *per se*, also support high richness of C4 grasses (Fig. 2b). C3 grass richness, on the other hand, is highest in the Northeast and the Trans-Himalaya (Fig. 2c).

The broad-scale relationships we observed between climate and other drivers on total, C3 and C4 grass species richness are in accordance with patterns reported in earlier studies from other regions (Bocksberger et al., 2016; Griffith et al., 2015; Hattersley, 1983; Liu et al., 2009; Teeri & Stowe, 1976; Visser et al., 2014). In general, species richness of all grasses, both C3 and C4, tended to increase with moisture index (Fig. 7c; (Bocksberger et al., 2016; Hattersley, 1983)), and decrease with precipitation seasonality (Fig. 7e). This increase in C4 richness with moisture index, which is somewhat counter to expectations, can be attributed to the fact that ∼75% of the C_4_ species in our dataset belong to the Panicoideae subfamily which is known to occur in warm and moist environments as compared to other subfamilies of PACMAD clade with C_4_ species (Bocksberger et al., 2016; Hartley, 1958; Lehmann et al., 2019; Taub, 2000; Visser et al., 2014). Further, as expected, C4 grass richness was greater in warmer areas, while C3 grasses tended to occupy cooler regions of the subcontinent (Fig. 7b, (Bocksberger et al., 2016; Bryceson & Morgan, 2022; Hattersley, 1983; Havrilla et al., 2023; Streit et al., 2024; Teeri & Stowe, 1976)). The preference of C4 species for warmer, drier conditions when compared to C3 species (Fig. 8) are as per expectations based on the physiological adaptations of C4 plants for photosynthesis at higher temperatures as compared to C3 (Ehleringer et al., 1997; Pau et al., 2013; Sage & Kubien, 2007). Finally, larger and more climatically heterogeneous districts (as indexed by MI and MI range) supported greater total and C4 grass richness (Fig. 7d, (Hattersley, 1983; Liu et al., 2009; Udy et al., 2021; Visser et al., 2014)), although this was not the case for C3 grasses. Studies on C3 grasses from other regions (Liu et al., 2009) have reported similarly weak relationships between area, heterogeneity and species richness. In our study, these patterns may have been more pronounced due to inconsistent relationships between district area and environmental heterogeneity in different regions and biogeographic zones of the subcontinent.

The richness and distribution patterns we observed for different subfamilies were also largely consistent with trends reported elsewhere (Bocksberger et al., 2016; Lehmann et al., 2019; Visser et al., 2014). The PACMAD clade consisting of both C3 and C4 species dominated in the warmer biogeographic zones, while the C3 subfamilies of Pooideae and Bambusoideae of the BEP clade dominated the cooler biogeographic zones of the country (Fig. 5a, Fig S1-S4; (Hartley, 1958; Lehmann et al., 2019; Visser et al., 2014)). Globally, Panicoideae which includes the Andropogoneae clade, is the most speciose clade, and is also known to be particularly abundant in the warm and wet tropics in areas with intermediate rainfall (∼1500mm; (Bocksberger et al., 2016; Hartley, 1958; Lehmann et al., 2019; Taub, 2000; Visser et al., 2014)). Panicoideae was also the most speciose subfamily in the Indian subcontinent, accounting for nearly half of all grass species in the country (Fig. 5), dominating all biogeographic zones, with its richness being highest in wetter, warmer and less seasonal districts (Fig. 5). Chloridoideae and Aristidoideae, on the other hand, were most speciose in warm and dry areas including the deserts and semi-arid biogeographic zones (Fig. 5), analogous to patterns reported elsewhere (Bocksberger et al., 2016; Edwards & Smith, 2010; Hartley, 1958; Lehmann et al., 2019; Liu et al., 2011; Taub, 2000; Valdés-Reyna et al., 2015).

The C3 subfamilies of Pooideae and Bambusoideae of the BEP clade had their greatest richness in the cooler biogeographic zones of the country, consistent with patterns reported in previous global and regional studies (Lehmann et al., 2019; Liu et al., 2011; Valdés-Reyna et al., 2015; Visser et al., 2014). The Pooideae subfamily, which tends to dominate temperate areas with cool and dry climates (Chapman, 1992; Lehmann et al., 2019), had the highest species richness in the Himalayas and the Trans-Himalayas (Fig. 5). The bamboos (subfamily Bambusoideae), on the other hand, were most speciose in the North-East which is a part of the Asia-Pacific hotspot of bamboo diversity (Fig. 5) (Bystriakova et al., 2003).

Our dataset, albeit at a coarse spatial scale, provides much-needed baseline information on the richness and distribution patterns of grasses of different photosynthetic types and subfamilies across the Indian subcontinent, as well as their underlying relationships with different climatic variables. Our findings have implications for the conservation of grasses and grasslands in the subcontinent, and can also serve as the basis for better understanding the potential responses of Indian grasslands to future climatic changes. Currently, much of the conservation effort in the country is focused on the well-known biodiversity hotspots of the Himalayas, North-East and the Western Ghats. However, our results suggest that other areas with high species richness of grasses, such as the arid Northwest and the Deccan Peninsula, also warrant conservation attention. Large tracts of these zones are covered by grassy biomes which are home to multiple endemics, specialised and threatened floral and faunal species. However, many of these areas have been misclassified as wastelands, and are under pressure from multiple drivers including land-use change, unscientific afforestation practices and invasive species (Kumar & Mathur, 2014; Mungi et al., 2020; Ratnam et al., 2016). Furthermore, most grass species in the country are range-restricted, with the Himalayas, Western Ghats and Deccan Peninsula in particular supporting a high number of such range restricted grass species which are potentially at risk from changes in climate and land use, necessitating greater conservation focus. Future efforts should focus on generating finer-scale datasets of grass species occurrences that also include abundance data, as it will allow for better distribution maps and a more nuanced understanding of richness-climate relationships and the potential impacts of future climate changes on the Poaceae of the Indian subcontinent.

## Supporting information

Supplementary_results

## Acknowledgements

We thank Arundhati Das and Sandeep Pulla for their help at different stages of the study. We thank National Centre for Biological Sciences, Tata Institute of Fundamental Research, Bengaluru for funding this study. Mukta Mande was awarded an intra-mural travel grant by National Centre for Biological Sciences to attend, and present the results of this study, in the 21^st^ Savanna Science Network Meeting, Kruger National Park, South Africa.

