## Supplementary_results for "Patterns of grass (Poaceae) species distribution and richness across India"

**Table S****1: List of Floras**

| **Sr. no.** | **Reference** | **Author** | **Publication Year** | **Publisher** |
| --- | --- | --- | --- | --- |
| 1 | A forest flora for the plains of Uttar Pradesh | P.C. Kanjilal | 1966 | Zucknow |
| 2 | A forest flora of the Andaman Islands | C.E. Parkinson | 1972 | International Book Distributors, Dehra Dun |
| 3 | Bengal Plants | David Prain | 1981 | Botanical Survey of India, Calcutta |
| 4 | Flora of Andhra Pradesh (India) | T. Pullaiah | 1997 | Scientific Publishers India |
| 5 | Flora of Assam | N.L. Bor | 1991 | Allied Book Centre, Dehra Dun |
| 6 | Flora of Bihar; analysis | N P Singh; V Mudgal; K K Khanna; S C Srivastava; A K Sahoo; S Bandopadhyay; N Aziz; M Das; R P Bhattacharya; P K Hajra | 2001 | Botanical Survey of India |
| 7 | Flora of Goa, Diu, Daman, Dadra and Nagar Haveli | Rolla Seshagiri Rao | 1986 | Botanical Survey of India |
| 8 | Flora of Great Nicobar Islands | B.K. Sinha (Auther)  P.K. Hajra & P.S. N. Rao (eds.) | 1999 | Botanical Survey of India |
| 9 | Flora of Gujarat State | Shah G.L | 1978 | K.A. Amin, Sardar Patel University Vallabh Vidyanagar |
| 10 | Flora of Haryana (Materials) | S. Kumar | 2001 | Bishen Singh Mahendra Pal Singh |
| 11 | Flora of Himachal Pradesh | H J Chowdhery, B M Wadhwa | 1984 | Botanical Survey of India |
| 12 | Flora of Jammu and Plants of neighbourhood | Brij Mohan Sharma & P. Kachroo | 1981 | Bishen Singh Mahendra Pal Singh |
| 13 | Flora of Karnataka; Analysis | B D Sharma, N P Singh, R S Raghavan, U R Deshpande | 1984 | Botanical Survey of India |
| 14 | Flora of Kerala - Grasses | P.V. Sreekumar & V.J. Nair | 1991 | Botanical Survey of India |
| 15 | Flora of Maharashtra State; Monocotyledones | B D Sharma, S Karthikeyan, N P Singh | 1996 | Botanical Survey of India |
| 16 | Flora of Pondicherry University | N. Parthasarathy, L. Arul Pragasan, C. Muthumperumal & M. Anbarashan | 2010 | Pondicherry University |
| 17 | Flora of Rajasthan | B.V. Shetty & V. Singh | 1993 | Botanical Survey of India |
| 18 | Flora of Saurashtra | P V Bole; J M Pathak | 1988 | Botanical Survey of India, Calcutta |
| 19 | Flora of Sikkim; Monocotyledons | P K Hazra, D M Verma, S Bandyopadhaya | 1996 | Botanical Survey of India |
| 2 | Flora of Tamil Nadu | A N Henry, V Chitra, N P Balakrishnan | 1989 | Botanical Survey of India |
| 21 | Flora of the Punjab plains | N.C. Nair | 1978 | Botanical Survey of India |
| 22 | Flora of Upper Liddar Valleys of Kashmir Himalaya | B.M. Sharma & P.S. Jamwal | 1998 | Scientific Publishers (India) |
| 23 | Forest flora of the Chakrata, Dehra dun and Saharanpur forest divisions, Uttar Pradesh | Upendranath Kanjilal & Basant Lal Gupta | 1969 | Manager of publications, Delhi |
| 24 | Grasses of Madhya Pradesh | G P Roy | 1984 | Botanical Survey of India |
| 25 | Grasses of North-Eastern India | U. Shukla | 1996 | Scientific Publishers |
| 26 | Plant diversity of Lakshadweep Islands | C. Sudhakar Reddy & P.S. Roy | 2011 | Bishen Singh Mahendra Pal Singh, Dehra Dun |
| 27 | Plants of Northern Gujarat | W T Saxton | 1978 | Bishen Singh Mahendra Pal Singh, Dehra Dun, India |
| 28 | Study of Angiospermic Flora of Kachchh District, Gujarat, India | Y S Patel, R M Patel, P N Joshi, Y B Dabgar | 2011 | Life sciences Leaflets |
| 29 | The flora of Delhi | J.K. Maheshwari | 1963 | Council of Scientific & Industrial Research, India |
| 30 | The flora of Orissa | H O Saxena, M Brahmam | 1996 | Orissa forest development corporation Ltd. |
| 31 | The flora of Tripura State | D.B. Deb | 2011 | Bishen Singh Mahendra Pal Singh, Dehra Dun |

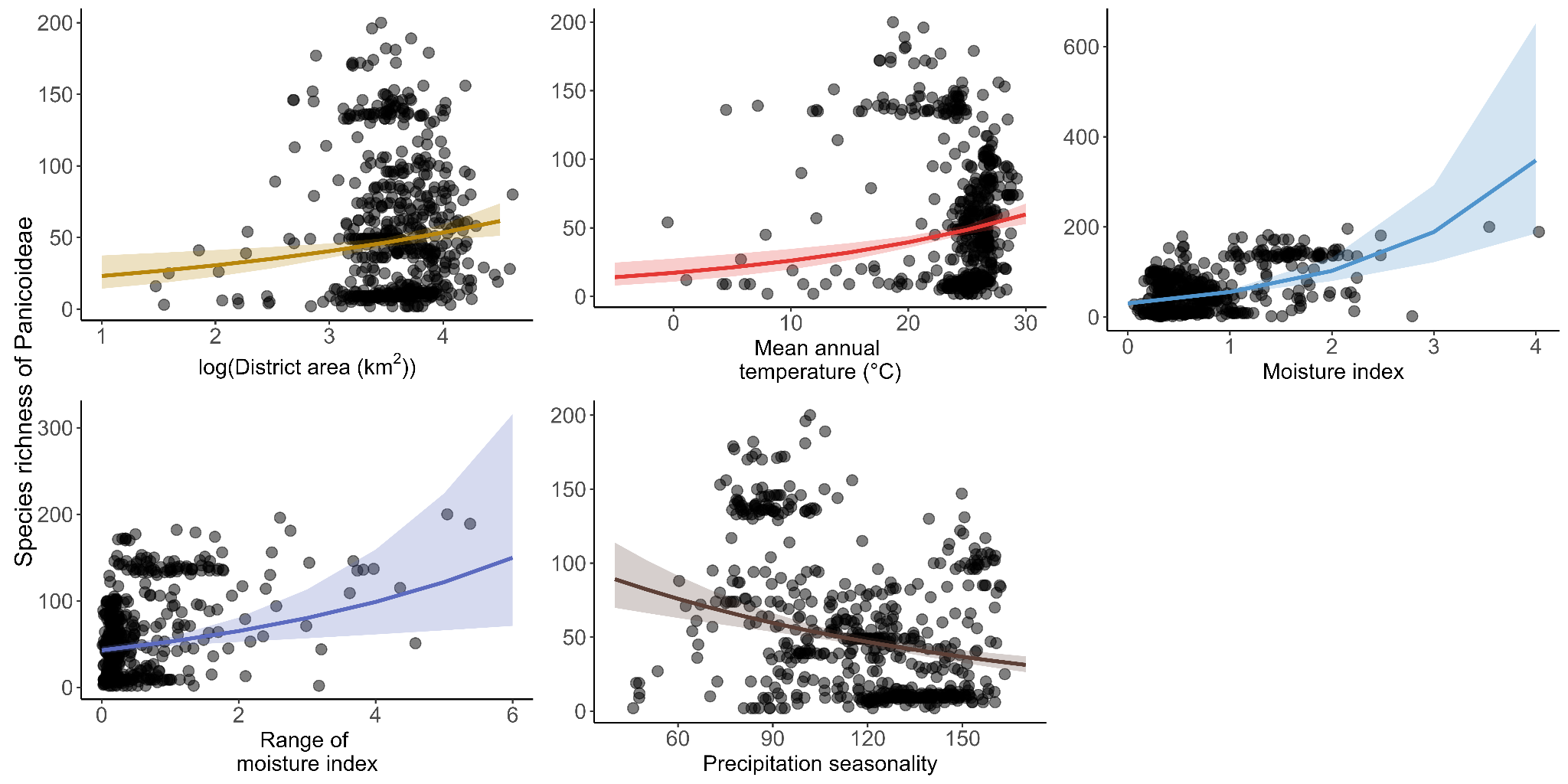

Fig. S1: Relationship between species richness of Panicoideae and different climatic variables (only the significant relationships are shown)

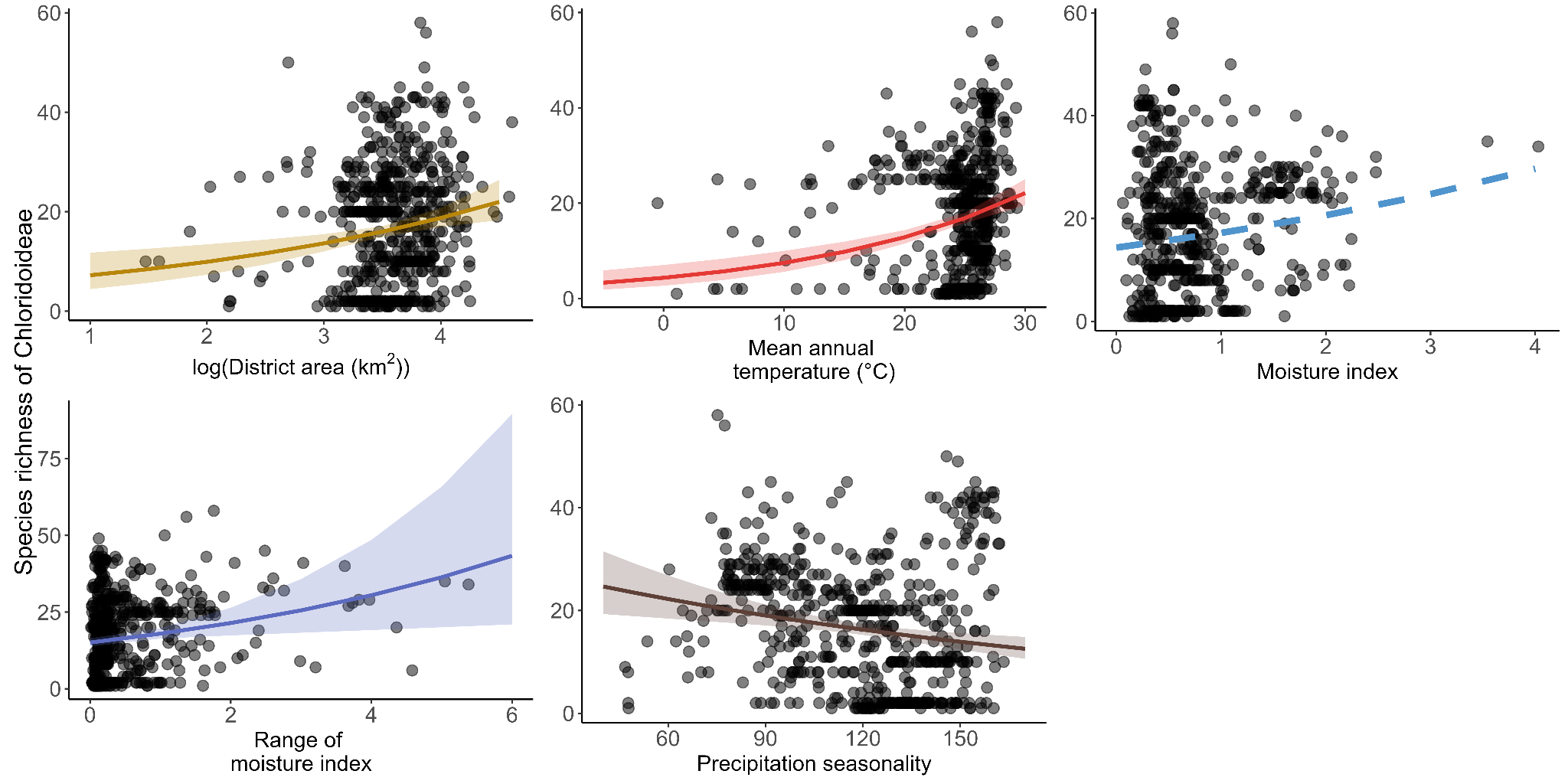

Fig. S2: Relationship between species richness of Chloridoideae and different climatic variables (only the significant relationships are shown)

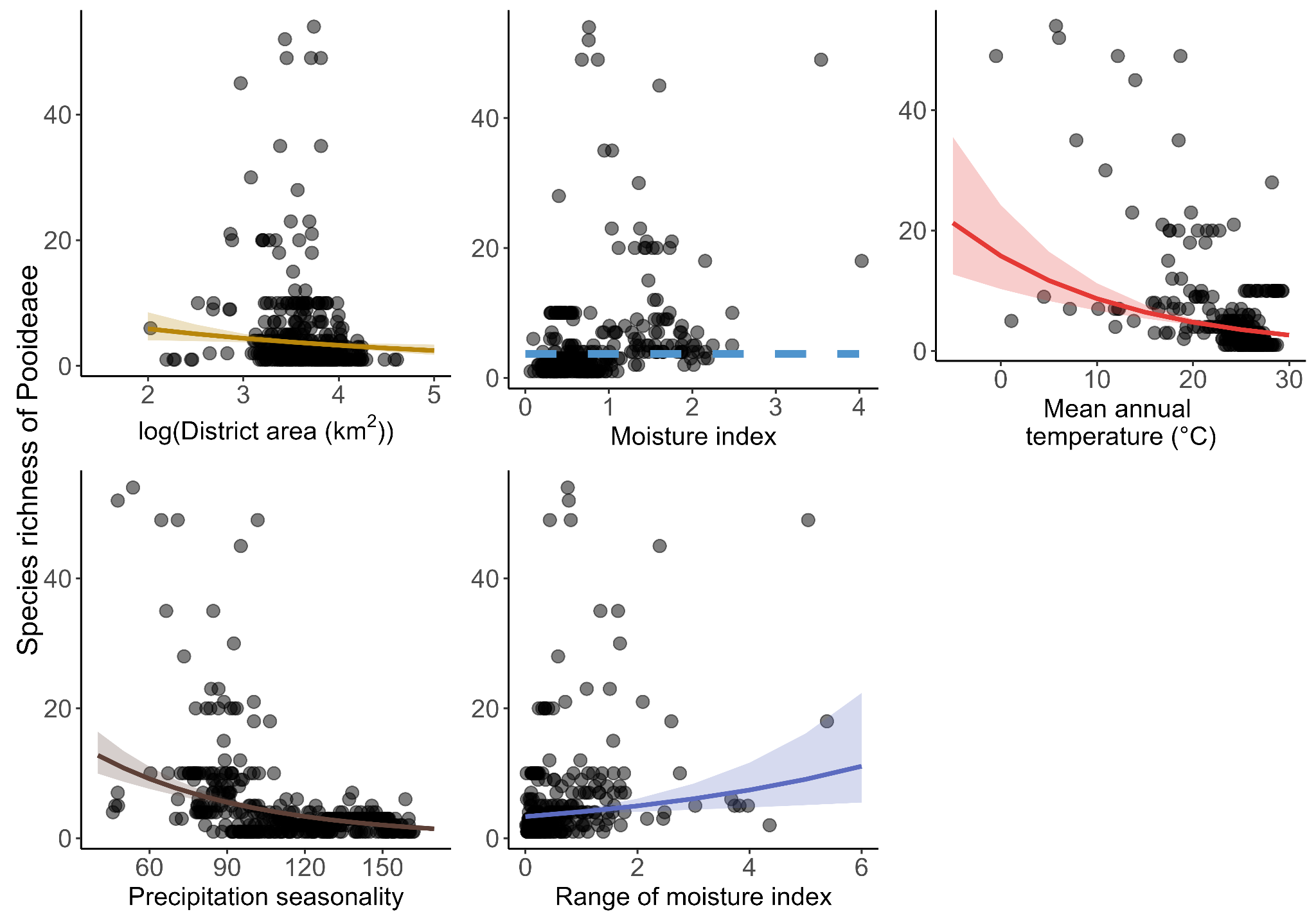

Fig. S3: Relationship between species richness of Pooideae and different climatic variables (only the significant relationships are shown)

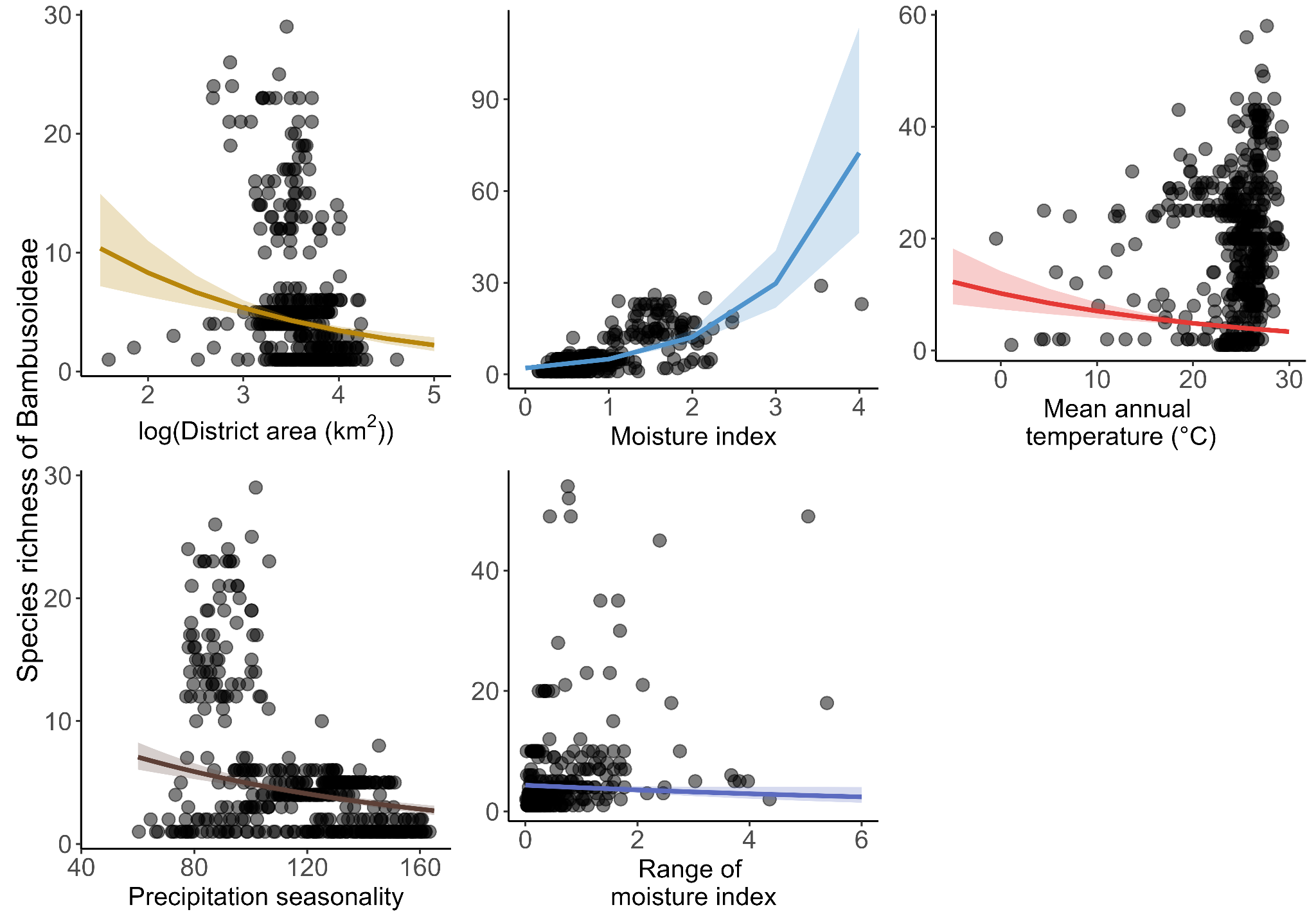

Fig. S4: Relationship between species richness of Bambusoideae and different climatic variables (only the significant relationships are shown)

**Table S2**

**AIC and R^2^ values for models with MAP vs AI as predictors. The models in bold indicate the models used for inference**

| ***Response*** | ***Predictors*** | ***AIC*** | ***pseudo-R^2^(Likelihood-ratio based)*** |
| --- | --- | --- | --- |
| Total species richness | log_10_(District area), MAP, MAT, PS, MAP range | 6354.452 | 0.242 |
| **Total species richness** | **log_10_(District area), AI, MAT, PS, AI range** | **6346.807** | **0.251** |
| Total species richness | log_10_(District area), AI, TW, PS, AI range | 6349.825 | 0261 |
| C_4_ species richness | log_10_(District area), MAP, MAT, PS, MAP range | 6019.397 | 0.183 |
| **C_4_ species richness** | **log_10_(District area), AI, MAT, PS, AI range** | **6015.655** | **0.188** |
| C_4_ species richness | log_10_(District area), AI, TW, PS, AI range | 6031.868 | 0.166 |
| C_3_ species richness | log_10_(District area), MAP, MAT, PS, MAP range | 3940.861 | 0.611 |
| C_3_ species richness | log_10_(District area), AI, MAT, PS, AI range | 3911.79 | 0.630 |
| **C_3_ species richness** | **log_10_(District area), AI, TW, PS, AI range** | **3889.538** | **0.643** |
| Proportion of C_4_ species | MAP, MAT, PS | 3302.368 | 0.902 |
| **Proportion of C_4_ species** | **AI, MAT, PS** | **3287.71** | **0.904** |
| Proportion of C_4_ species | AI, TW, PS | 3452.764 | 0.874 |

**Table S3: Model output for total species richness**

| ***Predictors*** | ***Incidence Rate Ratios*** | ***std. Error*** | ***CI*** | ***p*** |
| --- | --- | --- | --- | --- |
| **Intercept** | **14.83** | **6.10** | **6.49 – 34.57** | **<0.001** |
| **log(District area)** | **1.40** | **0.12** | **1.17 – 1.67** | **<0.001** |
| **Moisture index** | **1.76** | **0.17** | **1.47 – 2.11** | **<0.001** |
| **Mean annual temperature(°C)** | **1.03** | **0.01** | **1.02 – 1.05** | **<0.001** |
| **Precipitation seasonality** | **0.99** | **0.00** | **0.99 – 1.00** | **<0.001** |
| **Moisture index range** | **1.18** | **0.08** | **1.04 – 1.35** | **0.013** |
| **Observations** | **602** | | | |
| **R^2^ Nagelkerke** | **0.369** | | | |
| **AIC** | **6346.807** | | | |

**Table S4: Model output for C4 species richness**

| ***Predictors*** | ***Incidence Rate Ratios*** | ***std. Error*** | ***CI*** | ***p*** |
| --- | --- | --- | --- | --- |
| **Intercept** | **8.82** | **3.83** | **3.74 – 21.28** | **<0.001** |
| **log(District area)** | **1.36** | **0.13** | **1.12 – 1.63** | **0.001** |
| **Moisture index** | **1.58** | **0.15** | **1.31 – 1.91** | **<0.001** |
| **Mean annual temperature(°C)** | **1.05** | **0.01** | **1.03 – 1.07** | **<0.001** |
| **Precipitation seasonality** | **0.99** | **0.00** | **0.99 – 1.00** | **<0.001** |
| **Moisture index range** | **1.21** | **0.08** | **1.06 – 1.39** | **0.005** |
| **Observations** | **599** | | | |
| **R^2^ Nagelkerke** | **0.278** | | | |
| **AIC** | **6015.655** | | | |

**Table S5: Model output for C3 species richness**

| ***Predictors*** | ***Incidence Rate Ratios*** | ***std. Error*** | ***CI*** | ***p*** |
| --- | --- | --- | --- | --- |
| **Intercept** | **26.16** | **8.94** | **12.97 – 53.08** | **<0.001** |
| **log(District area)** | **1.17** | **0.09** | **1.00 – 1.37** | **0.043** |
| **Moisture index** | **2.41** | **0.17** | **2.10 – 2.78** | **<0.001** |
| **Maximum temperature of the wettest quarter(°C)** | **0.99** | **0.01** | **0.97 – 1.00** | **0.035** |
| **Precipitation seasonality** | **0.99** | **0.00** | **0.98 – 0.99** | **<0.001** |
| **Moisture index range** | **1.07** | **0.05** | **0.98 – 1.18** | **0.165** |
| **Observations** | **590** | | | |
| **R^2^ Nagelkerke** | **0.872** | | | |
| **AIC** | **3889.538** | | | |

**Table S6: Model output for proportion of C4 species**

| ***Predictors*** | ***Odds Ratios*** | ***std. Error*** | ***CI*** | ***p*** |
| --- | --- | --- | --- | --- |
| **Intercept** | **0.75** | **0.05** | **0.65 – 0.86** | **<0.001** |
| **Moisture index** | **0.78** | **0.01** | **0.76 – 0.81** | **<0.001** |
| **Mean annual temperature(°C)** | **1.04** | **0.00** | **1.04 – 1.05** | **<0.001** |
| **Precipitation seasonality** | **1.01** | **0.00** | **1.00 – 1.01** | **<0.001** |
| **Observations** | **599** | | | |
| **AIC** | **3287.710** | | | |
